# Designed nanofiber immunogens prime antibody responses to diverse conformations of the HIV-1 fusion peptide

**DOI:** 10.64898/2026.10.02.756244

**Authors:** Chelsea N. Fries, Jui-Lin Chen, Nicole L. Votaw, Benjamin Cossette, Emily F. Roe, Maria Dennis, Elizabeth J. Curvino, Luqman Issah, Kat M. Lazar, Shamitha Shetty, Rachel Katz, Corrine Nief, Sallie Permar, Koen K.A. Van Rompay, Genevieve Fouda, Joel H. Collier

## Abstract

Focusing antibody responses towards known neutralizing epitopes is a major strategy for protective immunization. Targeting flexible antigenic regions represents a unique challenge because functional antibodies must be able to bind the epitope of interest in multiple conformations. The fusion peptide (FP) of HIV-1 is a flexible peptide critical to viral entry of host cells for which broadly neutralizing antibodies have been identified. Because elicitation of potent, functional antibodies toward this epitope remains an unmet challenge in HIV-1 vaccination, we introduce a designed immunogen platform capable of raising anti-FP-antibodies with increased conformational tolerance relative to a traditional carrier protein. Antibodies induced by these immunogens bind heterologous FP sequences when elicited in mice and infant non-human primates. This platform shows promise for preclinical development to prime antibody responses toward FP of the HIV-1 envelope and can likely be translated to peptide immunofocusing for other viral pathogens.

## Introduction

Broadly neutralizing antibodies (bNabs) against HIV-1 most commonly target non-contiguous conformational epitopes that often include glycosylation sites, which has made selection of peptide epitopes for HIV vaccination challenging. However the fusion peptide (FP) of HIV-1 is the binding site of the bNab VRC34.01.^1^ FP is a relatively conserved 8-10 amino acid sequence at the envelope trimer’s N-terminus which can rotate and twist, assuming diverse conformations which allow HIV virions to infect host cells.^2,3^ The identification of this short, linear epitope provides an opportunity to generate peptide immunogens which could, in theory, generate protective antibody responses against HIV-1. However, immunization against FP often results in low-magnitude and low-affinity responses when it is displayed on scaffold proteins.^2,4–6^ The most effective method for inducing FP-directed antibody responses to date has been to prime the immune system with the carrier protein Keyhole Limpet Hemocyanin (KLH) decorated with FP peptides, followed by boosting with native-like trimers to mature antibodies towards FP presented on native HIV-1 virions.^5^ This type of heterologous immunization regimen has produced monoclonal antibodies with broad neutralization potential, but generation of anti-FP responses is extremely variable between individuals and requires boosting with SHIV to elicit neutralization.^5,7^ However, these studies show that fusion peptide is an excellent target and antibodies targeting it can provide broad and potent neutralization. On native HIV envelope trimers, FP assumes many different conformations.^2^ The structures of FP bound to unique bnAbs do not converge on a single stabilized FP conformation, but show that FP is flexible, and antibodies with many contact angles are capable of neutralizing the virus.^2^ Thus, priming immunogens which can produce antibody repertoires that can bind multiple conformations of FP are more likely to produce neutralizing FP-directed responses. For these reasons, improvement of FP-directed antibody responses has the potential to impact the generation of protective antibody responses to HIV-1.

Currently, there are very few immunogen platforms capable of generating robust antibody responses against flexible linear peptides. In preclinical settings, KLH has been the primary tool for priming antibody responses towards a specific peptide. KLH has several drawbacks for implementation, however, as it is prone to aggregation and difficult to express recombinantly with the motif of interest displayed on its exterior. Alternatively, using KLH relies on the ability to purify the antigenic peptide and conjugate it to the surface of the KLH carrier, a task that can be complicated for hydrophobic peptides. For these reasons, a platform which is capable of genetically fusing and displaying high copy numbers of flexible and/or hydrophobic peptides on a nanostructure would be highly advantageous for FP-directed immunization strategies and similar approaches.

To develop an immunogen platform for FP-directed immunization, we sought to tackle key challenges in existing FP-directed approaches. To focus responses on FP after heterologous immunization, FP-specific B cells must be sufficiently competitive against B cells reactive towards a vast number of other immunodominant epitopes on HIV trimers. In previous attempts to prime FP-specific responses, antibody specificity has drifted from FP toward other non-neutralizing epitopes upon boosting with SOSIP trimers, which is required for affinity maturation of FP antibodies.^8^ Systematic studies of germinal center dynamics have noted that increasing the number and affinity of precursor B cells within germinal centers can improve their competitive fitness upon boosting with antigens bearing multiple epitopes with varying levels of immunodominance.^9^ Building upon these findings, we hypothesized that improving the number and affinity of FP-reactive B cell clones with FP-priming immunogens would improve the fitness of these cells upon SOSIP boosting and increase the resultant seroprevalence of FP-directed antibodies.

A key pathway for improving the quality of responses primed towards FP is to increase the maturation of FP-directed antibodies within germinal centers (GCs). Increased accumulation of nanomaterials in germinal centers improves the number and affinity of B cells within GCs,^9,10^ an effect which is boosted by the decoration of nanostructures with glycans.^11^ The role of glycans in promoting material trafficking to GCs is restricted to materials of a certain size, however, as these materials must be small enough (<100 nm) to drain freely to lymph nodes to be transported to B cell follicles.^11^ Though these phenomena are known to improve B-cell responses, existing technologies have not utilized glycan-decorated nanomaterials to improve antibody responses against peptide epitopes. Previous studies have highlighted the ability of filamentous peptide nanostructures to raise antibody responses to peptide epitopes, with a particular capacity to generate responses towards epitopes with rare and low affinity B cell precursors.^12,13^

Coil29 is a coiled-coil peptide which self-assembles into filaments and contains multiple T cell epitopes, which generate strong T follicular helper responses in germinal centers, functional antibodies, and CD8^+^ T cell responses.^14–16^ Utilizing this modular and immunogenic platform, we combined tools which control nanofiber assembly and glycomimetic motifs to tune nanofiber trafficking properties. These immunogens were then employed to display HIV fusion peptides for raising antibodies towards FP which can bind FP in a variety of conformations.

## Results

### Design and Biophysical Characterization of Fusion Peptide Nanofiber Immunogens

To create immunogens with tunable size and glycomimetic character, we used multi-component assemblies containing up to 5 unique peptides. The nanofiber-forming peptide Coil29 has been widely used to display peptide epitopes and generate cellular and humoral immune responses without the need for additional adjuvants.^15–17^ Coil29 assembles in to coiled filaments in which opposite faces of α-helices pair to extend the nanofiber assemblies.^14^ These nanofibers typically extend in unbounded structures but can be stabilized and formed at defined sizes by incorporating Coil29Caps, designed peptides which limit assembly by creating a satisfied assembly edge to nanofiber structures.^18^ In this study, assemblies are made entirely from synthetic peptides including (a) Coil29 peptides to induce nanofiber assembly,^15,16^ (b) FP-Coil29 to display fusion peptide epitopes from HIV-1, (c) Glyco-Coil29 glycomimetic peptides to modulate immunogen trafficking,^19^ and (d) two Coil29Caps to control nanofiber length by binding to each end of the nanofiber.^18^ Each of these components can be added or removed from assembly formulations to test the impact of immunogen size and glycomimetic character, with the full formulation shown in Figure 1a (sequences in Table S1).

**Figure 1:**
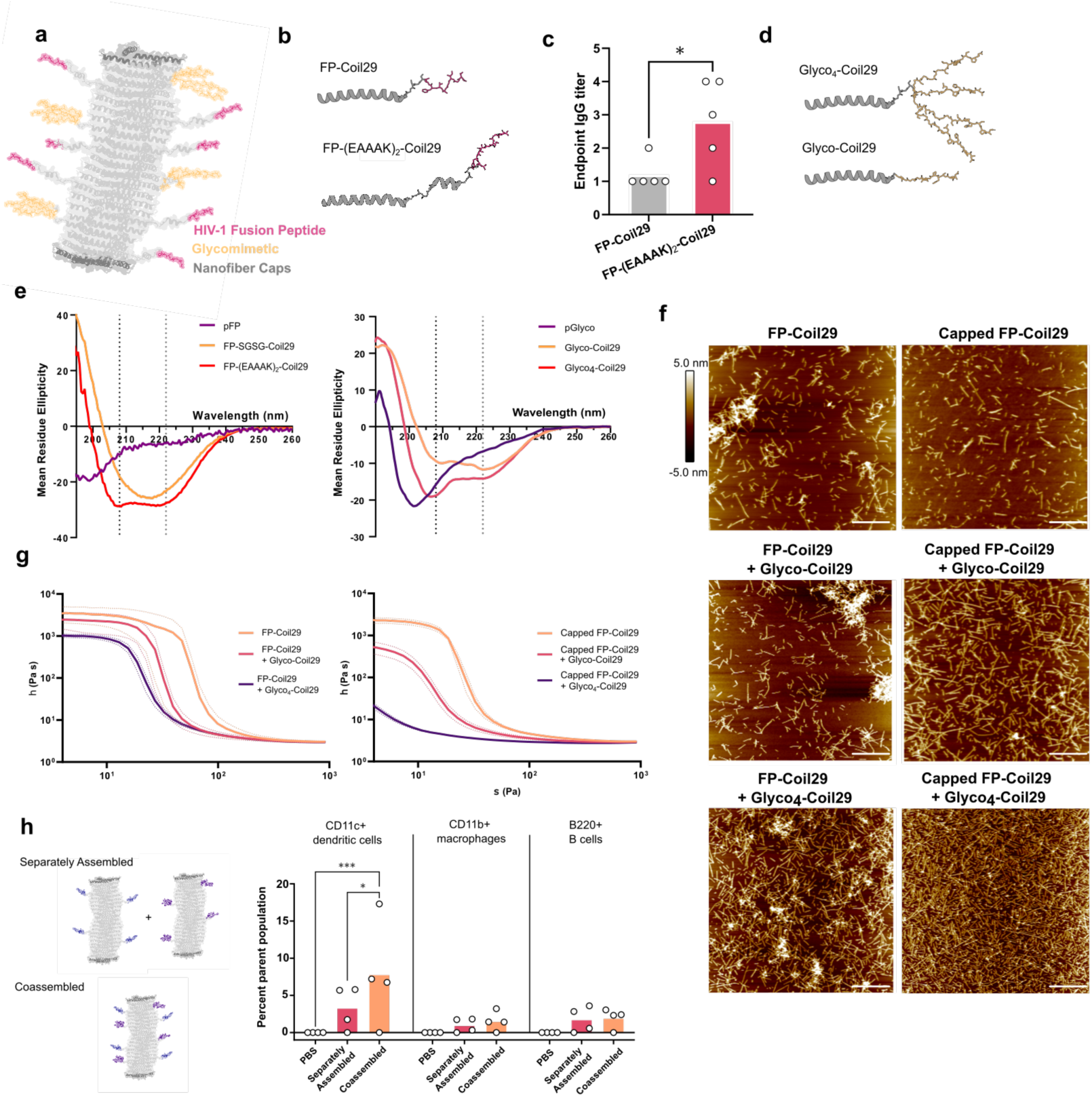
Design and biophysical characterization of Fusion Peptide nanofiber immunogens. **(a)** Schematic of FP-Coil29 nanofibers based on the structure of Coil29 (PDB 3j89) capped with size-controlling Coil29Caps (dark grey), FP fusions (pink) and glycomimetic peptides (gold). **(b)** Structure of FP-Coil29 peptides with helical Coil29 motif and helical EAAAK spacers (grey) with N-terminally linked FP sequence AVIGIAVFL (pink). **(c)** Antibody titers of C57BL/6 mice immunized with FP-Coil29 with or without helical EAAAK spacers. Mice were immunized at 0, 2, and 4 weeks. IgG titers to FP were measured at 5 weeks after primary immunization. Animals immunized with FP-(EAAAK)_2_-Coil29 constructs showed significantly higher anti-FP IgG titers (p=0.04, unpaired t-test with Welch’s correction). **(d)** Glycomimetic peptides (gold) displayed adjacent to the Coil29 sequence (grey) with either a linear peptide linker (Glyco-Coil29) or after branching lysine resides (Glyco_4_-Coil29). **(e)** Circular Dichroism of unmodified fusion peptide, and fusion peptide linked to Coil29 via a SGSG- or EAAAK-linker (left). Circular Dichroism of soluble glycomimetic peptides and its fusions to Coil29 in linear monomeric and branched tetrameric format (right). **(f)** AFM images of nanofiber formulations assembled in PBS. Images displayed above are representative height maps. (scale bar = 800 nm) **(g)** Viscosity of FP-Coil29 nanofibers formulated with capping and glycomimetic peptides was measured using a viscometry shear stress ramp from 0.1 to 1000 Hz**. (h)** Uptake of co-assembled and separately-assembled linear and branched peptides by immune cells in lavage fluid after intraperitoneal injection compared by one-way ANOVA with Fisher’s LSD post-hoc analysis (*p<0.05, ***p<0.001).

Our first goal was to design an immunogen with predictable structure, solubility, and capacity to induce FP-specific antibodies. To achieve this, we generated constructs which displayed the Fusion Peptide of HIV-1 (AVGIGAVFL) linked to Coil29. Coil29 nanofibers assembled well when a rigid spacer composed of two repeats a rigid helical linker (EAAAK) was used to extend hydrophobic FP epitopes away from Coil29 assembly motifs (Figure 1b). Conversely, constructs which use a short SGSG linker generated extremely stiff, hydrogel-like solutions which were prone to precipitating over time. We also expected that the helical spacer would increase the availability of FP epitopes for binding on the surface of B cells. Indeed, when we immunized mice with both FP-Coil29 and FP-(EAAAK)_2_-Coil29 constructs, mice raised significantly higher antibody titers when the FP epitope was extended from the Coil29 motif (Figure 1c). After establishing a baseline for Coil29 nanofibers displaying FP, we incorporated glycomimetic nanofiber components which aimed to maximize lymph node accumulation and trafficking to germinal centers within them.

To modulate trafficking to GCs, we employed peptides based on a sequence that was predicted *in silico* and experimentally validated for binding to various lectins^20^ including CD169,^19^ a receptor expressed on macrophages which transport particulate materials to B cell follicles.^21^ In addition to the complement-mediated transport of glycosylated materials mentioned above, we hypothesized that increasing association of our materials with CD169^+^ macrophages could alter the affinity maturation of B cells encountering immunogens via this pathway. We synthesized tetrafunctional peptide variants of this sequence by introducing branching lysines after the linear Coil29 sequence in addition to synthesizing a monofunctional variant (Figure 1d).

We employed the Coil29 backbone to assemble the FP epitope and glycomimetic peptides into defined nanostructures and generated formulations with or without capping peptides. When measured by circular dichroism, both FP and Glyco peptides are relatively unstructured on their own, but when appended to Coil29 show an α-helical secondary structure, indicating the Coil29 motif retained helicity after fusing FP and both linear and branched displays of Glyco sequences (Figure 1e). Each of these formulations formed nanofibers as imaged by AFM (Figure 1f). Notably, the nanofibers aggregated more readily in samples without capping peptides. This presence of nanofiber aggregates noted by AFM was subsequently confirmed by viscometry (Figure 1g). In nanofiber formulations without capping peptides, the inclusion of glycomimetic peptides did not significantly alter properties. In capped formulations, however, glycomimetic peptides reduced the viscosity of nanofibers (Capped FP-Coil29 + Glyco-Coil29) and produced markedly less viscous nanofibers when incorporated in a branched format (Capped FP-Coil29 + Glyco_4_-Coil29). We also observed a phenomenon in which capping peptides generated slightly longer nanofibers than uncapped formulations containing Glyco-Coil29 or Glyco_4_-Coil29 (Figure S1). In contrast to the use of Coil29Caps to shorten nanofibers which persist to unbounded lengths when displaying other epitopes,^22^ they functioned as stabilizing agents in this system (further discussion in supplementary text).

We also used cellular uptake *in vivo* as a proxy for measuring the extent to which linear or branched peptides co-assembled into mosaic nanostructures using a previously described method.^17^ We appended TAMRA to Coil29 and Alexafluor647 to Coil29 after introducing a tetrafunctional branching point. We then constructed nanofiber assemblies where these two peptides were mixed prior to assembly (co-assembled) or nanofibers were formed separately and mixed immediately before intraperitoneal injection in mice (separately assembled). After 4 hours, mice were sacrificed and lavage fluid was harvested to measure immune cells which had internalized fluorescently labeled nanofiber assemblies. We observed cells positive for both TAMRA and Alexa647 in both co-assembled and separately assembled formulations, and a greater number of double positive CD11c+ dendritic cells in mice injected with co-assembled formulations (Figure 1h, Figure S3,S4). This indicated that even in the case of separately assembled nanofibers, immune cells internalize nanofibers containing both linear and branched peptides. Furthermore, this effect is enhanced by mixing peptides prior to solubilizing them and triggering assembly with physiological salts.

### Trafficking of size-controlled, glycomimetic immunogens to lymph nodes boosts germinal center responses

Nanomaterial size is a well-appreciated factor which dictates lymphatic drainage of immunogens, but the effect of solution-phase properties such as viscosity of fibrillar materials on lymph node (LN) accumulation has not been thoroughly explored. Because we observed stark differences in the solution properties of FP-Coil29 nanofibers with capping and glycomimetic peptides, we hypothesized that these formulations would also behave differently as they drain through extracellular space and flow in lymphatic vessels. Likewise, immunogen capture and accumulation are influenced by glycosylation, so we expected that glycomimetic peptide content would also play a role in the phenomenon. To test the effect of each of these factors on filamentous materials, we incorporated Coil29 peptides labeled with AlexaFluor647 into nanofiber assemblies and measured their accumulation in draining lymph nodes after subcutaneous injection. Inguinal lymph nodes were excised at 6, 12 and 48 hours after tail base injection and measured via IVIS to quantify the amount of material present in each node (Figure 2a,b, FigureS4).

**Figure 2:**
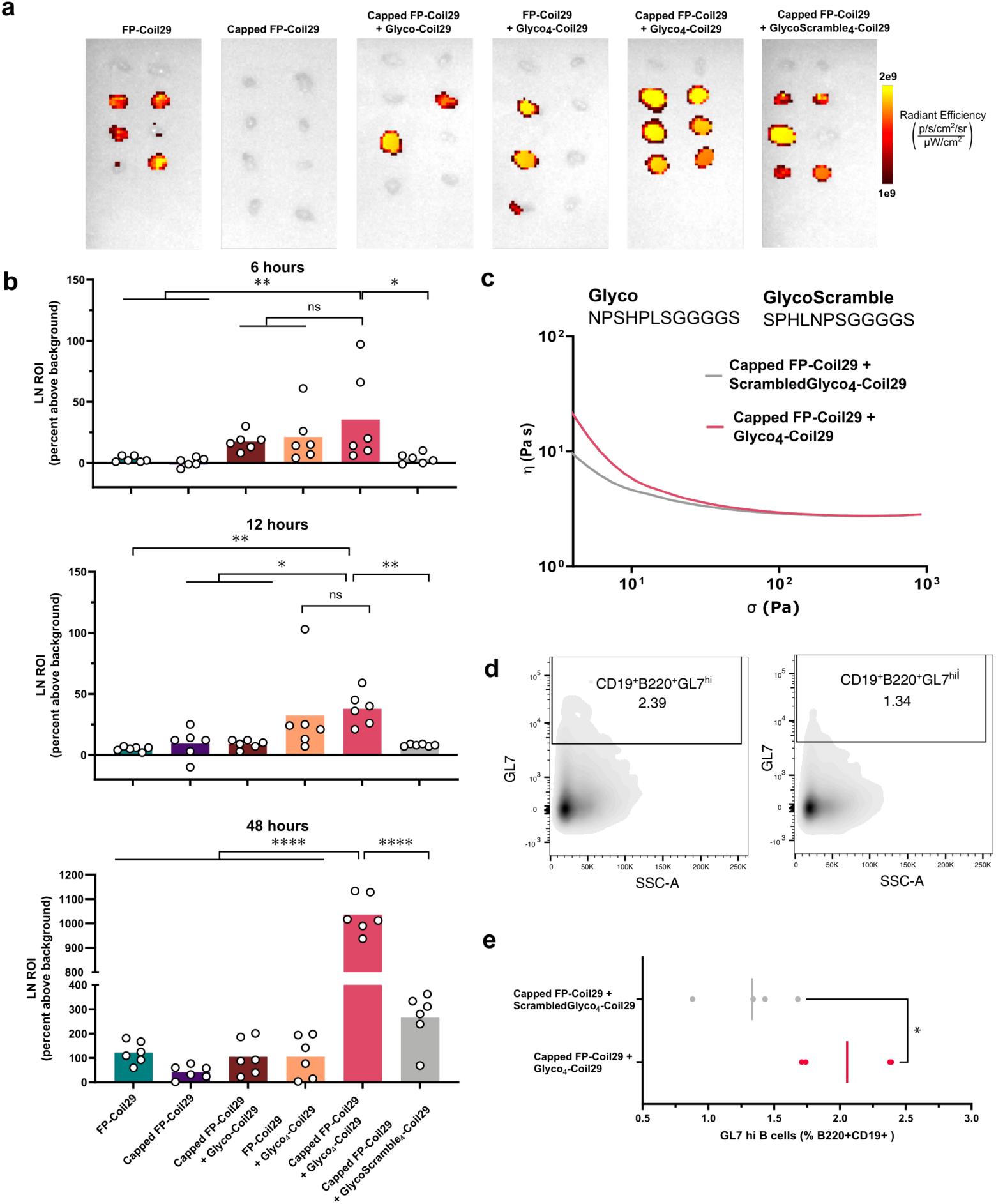
Nanofiber trafficking to draining lymph nodes and germinal center responses to glycomimetic immunogens. **(a)** Inguinal lymph nodes which drain directly from tail base subcutaneous injection sites were excised at 48 hours post-injection and imaged by IVIS to detect Alexa647-labeled peptide assemblies. **(b)** Quantification of radiant efficiency in lymph nodes relative to naïve mice after ex vivo imaging by IVIS at 6-, 12-, and 48-hours after injection compared to untreated mice (top row) (Images from 6- and 12-hours shown in Figure S4). Groups were compared by Ordinary One-Way ANOVA at each time point with Dunnet’s multiple comparison testing for pairwise comparisons (*p<0.05, *p<0.01, ***p<0.001, ****p<0.0001). **(c)** Viscosity of Capped FP-Coil29 nanofibers formulated scrambled and unscrambled glycomimetic peptides was measured using a viscometry shear stress ramp from 0.1 to 1000 Hz. **(d)** Mice were immunized via subcutaneous tail base injection 3 times in 2-week intervals and sacrificed 1 week after the final immunization. Draining inguinal lymph nodes were harvested and processed for flow cytometry to detect B220^+^CD19^+^GL7^hi^ germinal center B cells. **(e)** Percentage B220^+^CD19^+^ cells gated for high GL7 expression (gating shown in Figure S5) were compared using a Mann Whitney non-parametric test (*p<0.05).

We detected fluorescence earlier in the lymph nodes of mice injected with nanofibers containing linear and branched glycomimetic peptides regardless of capping peptide content, suggesting that the glycomimetic peptide sequence promoted retention of nanofibers in lymph nodes after passive drainage. After 48 hours, when both active and passive transport play a role in nanomaterial trafficking, we detected a striking increase in material for Capped FP-Coil29 + Glyco_4_-Coil29 relative to all other nanofiber formulations. While the full mechanism driving these differences are not clear, the results suggest that both glycomimetic peptide content and unaggregated-nanofiber solution properties are important for promoting lymphatic trafficking and retention (Figure 2a,b).

Because we observed differences in the biophysical properties of Capped glycomimetic nanofibers, we asked whether the reduction of nanofiber aggregation was responsible for improved lymph node accumulation, or if this effect was dependent on the glycomimetic peptide sequence we selected for this study. To test this, we generated a branched peptide with a scrambled glycomimetic sequence (Figure 2c). We expect the scrambled peptide to have similar physicochemical properties without the glycomimetic properties. Indeed, although Capped nanofibers made with scrambled and unscrambled glycomimetic sequences exhibited similar viscometry profiles, the designed glycomimetic sequence showed a strong preference for lymph node accumulation compared to its scrambled comparison as measured by IVIS (Figure 2a,b), indicating that the sequence-specific properties of this motif are key to its activity. We also immunized C57BL/6 mice with 3 doses of capped nanofibers incorporating branched peptides with a lectin-binding or scrambled glycomimetic sequence. After the final immunization, the draining lymph nodes of mice which received the Glyco_4_-Coil29 formulation produced significantly more germinal center B cells than scrambled (GlycoScramble_4_-Coil29) immunizations (Figure 2d,e). Because of the improved trafficking and germinal center responses we observed for Capped FP-Coil29 + Glyco_4_-Coil29 immunizations, we analyzed their activity as immunogens.

### Size-controlled glycomimetic immunogens raise antibody responses to multiple conformations of HIV-1 fusion peptide

Because of the high-density display afforded by incorporating branched peptides into assemblies, we measured antibody responses to the glycomimetic sequence on Glyco_4_-Coil29 when this peptide was added into assemblies at varying stoichiometries. After 3 immunizations, mice developed antibodies against the Glyco sequence when nanomaterials contained 25% or 10% Glyco_4_-Coil29 (Figure 3a), but not when Glyco_4_-Coil29 was excluded from formulations. When immunized with 25% Glyco_4_-Coil29, FP-specific antibodies were diminished, but in mice immunized with 10% Glyco_4_-Coil29, FP-specific antibodies were induced at a level not statistically different from FP-Coil29 alone. This suggests that branched peptides may sterically occlude FP peptides when included at a high density in assemblies, or anti-Glyco responses may compete with anti-FP responses, though this phenomenon is dependent on the relative immunodominance of those two epitopes.^23^ Because 10% Glyco_4_-Coil29 elicited robust levels of anti-FP antibodies, this formulation was used in subsequent studies.

**Figure 3:**
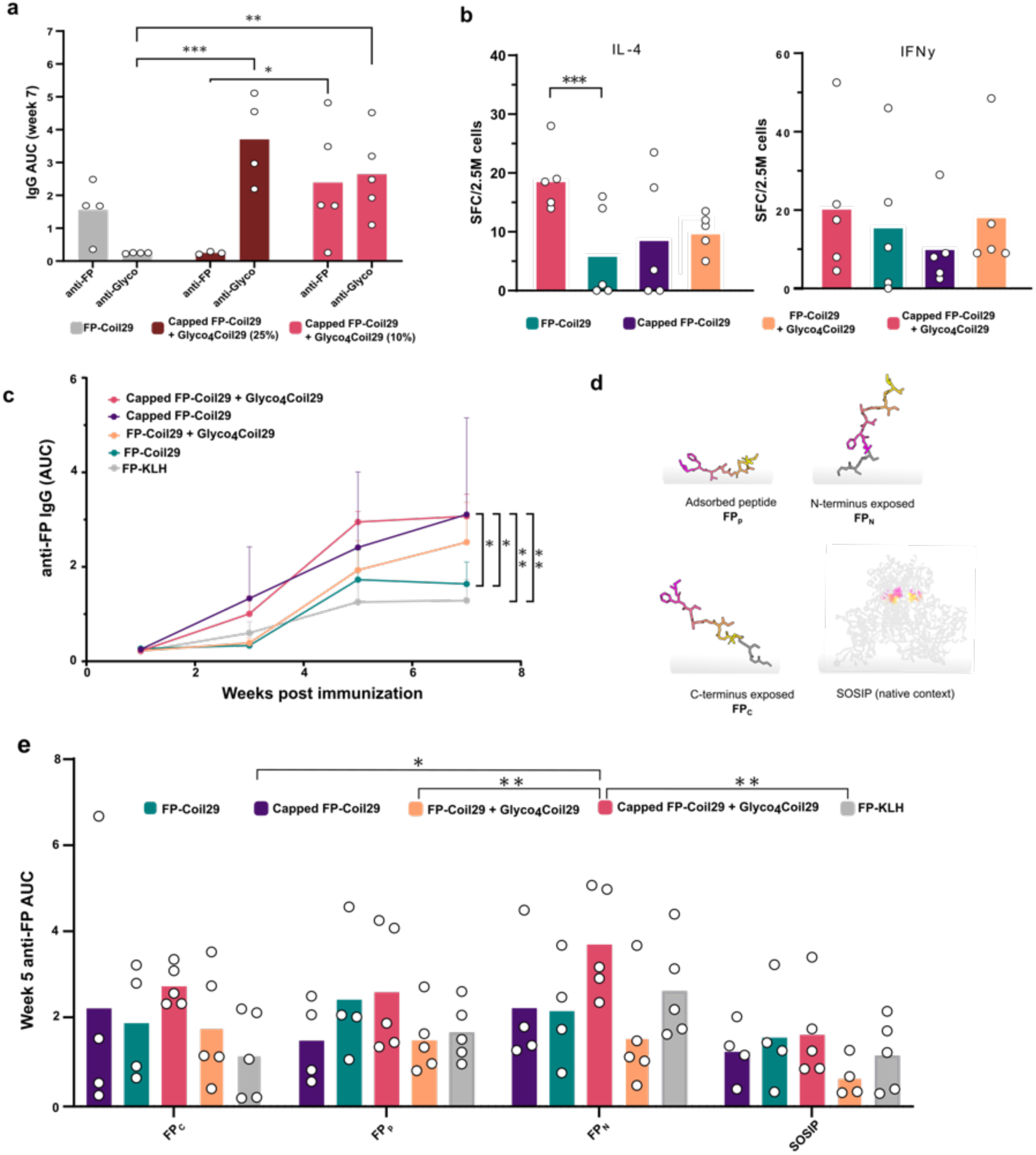
Antibody Responses to Unadjuvanted Fusion Peptide Immunogens. In all studies, mice were immunized via subcutaneous tail base injection 3 times in 2-week intervals and sacrificed 1 week after the final immunization. **(a)** FP-Coil29 immunizations with 10% or 25% Glyco_4_-Coil29 were given at 0, 2, and 4 weeks and blood was sampled from mice at 7 weeks after the first immunization. Antibody binding to FP and Glyco peptides was measured by ELISA. Antibody responses to each immunization were compared using 2-way ANOVA and responses to FP and Glyco peptides were compared across groups using Fisher’s LSD test. **(b)** Splenic lymphocytes were harvested and stimulated with the Coil29 peptides. After 48 hours of incubation, IL-4 and IFNγ producing lymphocytes were detected by ELISPOT. Groups were compared 1-way ANOVA and Fisher’s LSD test for multiple comparisons. **(c)** IgG binding to FP_N_ was measured over the course of the regimen and responses were compared between groups using repeated measures ANOVA. **(d)** Schematic representation of antigenic probes bound to ELISA plates. **(e)** IgG responses to diverse FP probes from sera in (c). Differences between groups were compared using two-way ANOVA (p=0.014 difference between treatment groups, p=0.008 difference between binding probes) and differences between treatment groups were subsequently compared using Fisher’s LSD test. *p<0.05, **p<0.01 ***p<0.005

We analyzed T cell responses to FP-Coil29 immunogens to determine if LN accumulation also impacted the extent of T cell help induced by these formulations. After 3 immunizations, spleens were harvested from immunized mice and stimulated with immunizing peptides for analysis by ELISPOT. Coil29 contains multiple T cell epitopes which increase its immunogenicity in mice^16^ and Coil29 reactive T cells were detected in the splenocytes of all mice immunized with FP-Coil29 nanofibers. We observed an increase in Coil29-reactive IL-4 producing T cells for mice immunized with Capped FP-Coil29 + Glyco_4_-Coil29, which is consistent with the increased nanofiber drainage to the lymph nodes of these mice (Figure 3b).

To test the effect of nanofiber capping and glycomimetic content on antibody responses, mice were immunized with FP-Coil29 nanofibers including either capping peptides, glycomimetic peptides, or both and compared to the peptide immunogen carrier KLH. The KLH variant used in this study utilized a rigid helical spacer (EAAAK) to display FP, which matched nanofiber displays, but differs from previously published variants which have utilized a Cysteine-maleimide linkage without additional spacer sequences.^6,24^ After immunization, all Coil29-scaffolded FP immunogens produced FP-reactive antibodies (Figure 3c). Over the course of the immunization regimen, Capped FP-Coil29 + Glyco_4_-Coil29 and Capped FP-Coil29 induced higher levels of FP-reactive antibodies than FP-Coil29 and FP-KLH constructs.

Because the ability of antibodies to bind conformationally distinct FP variants is key for generating neutralizing responses,^2^ we assessed antibody binding to FP via a variety of ELISA techniques in which the antigen was immobilized onto the plate in different conformations (schematics in Figure 3d). Specifically, FP was bound to ELISA plates with an available N-terminus (FP_N_), available C-terminus (FP_C_) or unmodified FP peptide directly adsorbed to ELISA plates (FP_p_) (all sequences and modifying chemistries described in Methods section). Additionally, SOSIP trimers were used to probe antibody binding to FP in a native-like conformation.

FP_N_ displays the FP sequence (AVGIGAVFL) in the same N-terminal format as in Coil29 and KLH immunogens and is extended from the surface of ELISA plates via a PEG linker. Likely because this probe matched the display of the epitopes in immunogens and is most sterically accessible, it showed the highest levels of binding in all immunization groups with the most consistent responses across individual animals when compared to the other probes tested. Binding to FP_C_ and FP_p_ was more variable and occurred at lower levels, possibly because antibody binding to these epitopes may require more unique angles of approach as these peptide conformations are less solvent-available than FP_N_. Finally, SOSIP trimers captured by apex-binding PGT145 antibodies displayed the FP epitope in its native-like conformation. Across these coating conditions, Capped FP-Coil29 + Glyco_4_-Coil29 induced a higher magnitude of antibodies which bound the diverse FP conformations displayed by each of these ELISA methods relative to Capped FP-Coil29, FP-Coil29 + Glyco_4_-Coil29, and FP-KLH (Figure 3e). Because Capped FP-Coil29 + Glyco_4_-Coil29 immunizations induced higher magnitude of anti-FP antibodies and these antibodies bound FP in a broader profile of display contexts, we advanced to studying them as priming immunogens for priming responses to HIV trimers.

### Priming immunogens shape antibody binding profiles after boosting with native-like HIV envelope trimers

To generate epitope-focused responses, immunogens are typically delivered sequentially, first by priming with a reductionistic design that presents target epitopes in the absence of the other competing sites in the native antigen, then subsequently boosted with native-like presentations of the full antigen to mature antibody responses toward conformations displayed on intact pathogens.^4,25^ To test the efficacy of FP-Coil29 nanofibers in this context, we administered 3 doses of FP-Coil29 immunogens formulated with CpG adjuvant to mice followed by 3 doses of SOSIP trimer, also formulated with CpG. Throughout this regimen, we monitored antibody responses to SOSIP, FP_N_, FP_C_ and FP_p_.

After 3 doses of these immunogens, all mice developed antibodies towards FP_N_ which continued to expand after SOSIP boosting (Figure 4a), indicating that the FP B cell lineage developed by priming immunogens was also reactive to SOSIP trimers. As this regimen progressed, the magnitude of FP antibodies induced by Capped FP-Coil29, Capped FP-Coil29 + Glyco_4_-Coil29, and FP-KLH were greater than those induced by FP-Coil29 or FP-Coil29 + Glyco_4_-Coil29.

**Figure 4:**
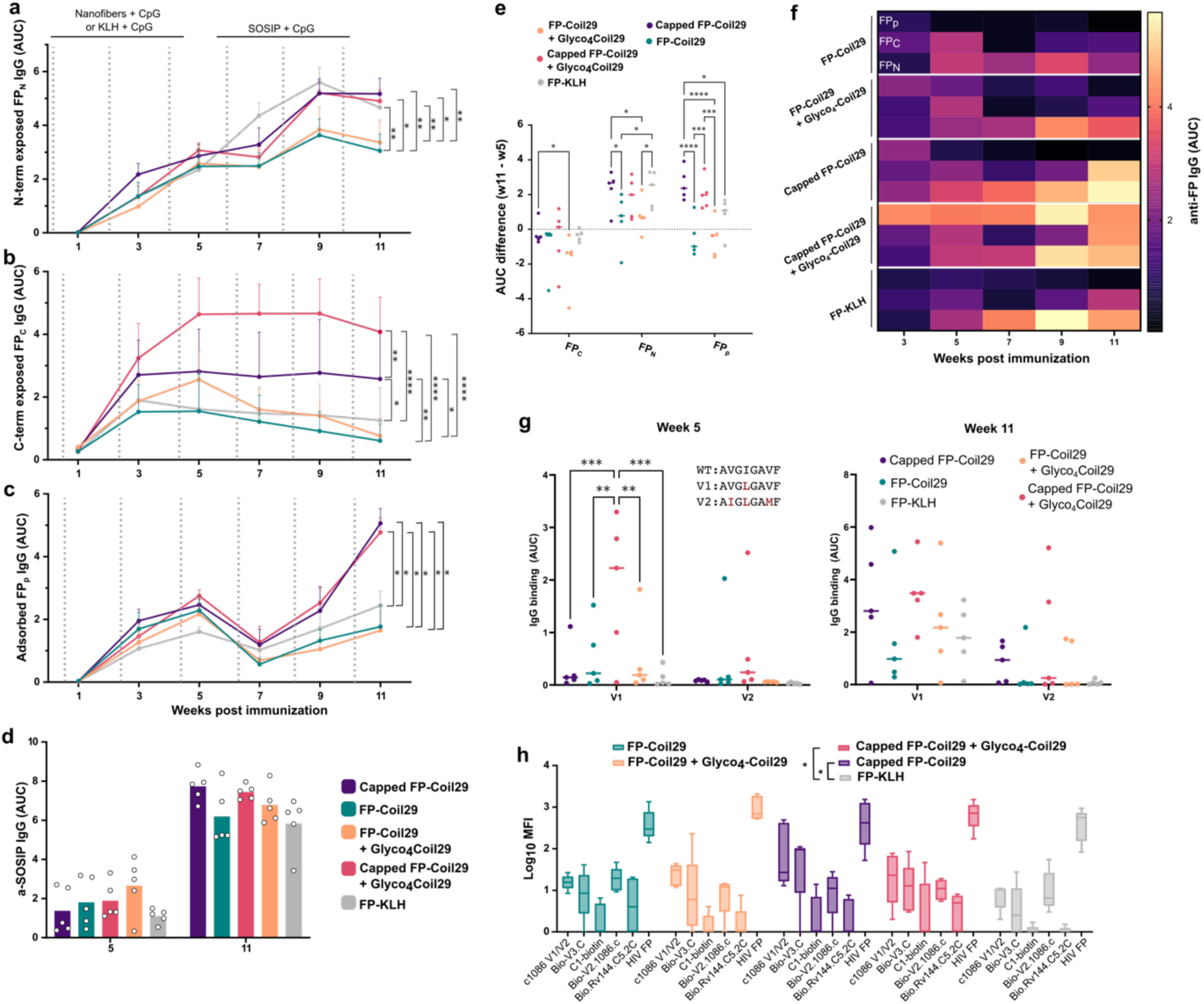
Fusion Peptide Immunogens prime distinct antibody profiles in sequential immunization regimens. **(a)** Serum IgG binding to FP_N_ measured by ELISA from biweekly blood collected from mice immunized as shown by arrows above graph. **(b)** Serum IgG binding to FP_C_ measured by ELISA from biweekly blood collected from mice immunized as shown by arrows above graph. **(c)** Serum IgG binding to FP_p_ measured by ELISA from biweekly blood collected from mice immunized as shown by arrows above graph. **(d)** SOSIP binding antibodies were measured after FP priming (week 5) and after multiple doses of SOSIP (week 11). For a-d, groups were compared using repeated measures one-way ANOVA and Fisher’s LSD post-hoc analysis. **(e)** Change in IgG responses to FP display formats after boosting with SOSIP (week 11 AUC relative to week 5 AUC). Groups were compared using 2-way ANOVA and multiple comparisons were made between experimental groups for each FP display format using Fisher’s LSD post-hoc analysis. **(f)** Heat map representation of mean antibody titers to FP_N_, FP_C_, and FP_p_ ELISA probes after immunization with FP immunogens. Responses to each probe are labeled for the first group and the pattern repeats across the lower rows. **(g)** IgG binding to heterologous FP sequences from commonly circulating strains. Binding was measured at week 5 (left) and week 11 (right). For each time point, binding to FPv1 and FPv2 were compared between groups using repeated-measures 2-way ANOVA and individual comparisons were subsequently made using Fisher’s LSD test. **(h)** Multiplexed IgG binding to a panel of epitopes from HIV Env after heterologous prime-boost with FP immunogens and SOSIP. Treatment groups were compared using repeated-measures 2-way ANOVA and individual groups were subsequently compared across all epitope specificities using Fisher’s LSD test.

The levels of FP-binding antibodies over the course of the FP SOSIP regimen showed differential evolution as we measured antibody binding to FP_C_ and FP_p_. Particularly for FP_C_ antibodies, SOSIP boosting caused the decay of this antibody subset upon boosting, indicating that antibody responses evolved toward a particular conformation which was not displayed by FP_C_ (Figure 4b,f) . However, mice which were primed with Capped FP-Coil29 + Glyco_4_-Coil29 retained high levels of FP_C_ binding antibodies, indicating that B cells which recognize this FP conformation remained competitive after multiple boosts. Mice primed with Capped FP-Coil29 also showed persistence of FP_C_ reactive antibodies, indicating the B cell responses induced by capped nanofiber immunogens develop unique binding properties compared to uncapped nanofibers and KLH. Similarly, we observed differences in antibody binding to FP_p_, wherein capped nanofibers induced stronger antibodies to this FP conformation over the course of the FP SOSIP immunization regimen (Figure 4c). Although no single ELISA method is indicative of superior FP antibody functionality, generating an antibody response which can bind structurally diverse conformations of FP is desirable for generating neutralizing responses to FP.^2^

The antibodies produced after FP-directed priming also bound SOSIP by ELISA (Figure 4d), though differences in SOSIP binding were not statistically different between priming immunogens. SOSIP-reactive antibodies continued to expand throughout the immunization regimen and likely included many epitope specificities after 3 doses of SOSIP were administered.

To quantify the change in binding to a given FP conformation after SOSIP boosting, we compared the difference in binding after 3 SOSIP boosts relative to binding after 3 doses of FP immunogens (Figure 4e). The mean change was positive across all 3 conditions only for mice immunized with Capped FP-Coil29 + Glyco_4_-Coil29. Mice primed with FP-KLH showed expansion of FP_N_ binding antibodies, but responses toward FP_C_ and FP_p_ decayed. The largest range of changes occurred for binding to FP_p_. Antibody responses expanded to a greater extent for animals immunized with capped nanofibers relative to FP-KLH and diminished responses from animals immunized with uncapped nanofibers. The combination of these antibody responses can also be visualized in parallel using the heatmap (Figure 4f) where each row displays the antibody response to a given coating over the course of the regimen for mice primed with FP-Coil29 or KLH immunogens.

Beyond measuring the binding of antibodies to FP in different display formats, we asked whether these sera were capable of binding FP sequences which are mutated from the immunizing sequence. FPv1 (AVG<u>L</u>GAVF) and FPv2 (AIG<u>L</u>GA<u>M</u>F) are the most common FP sequences found in circulating variants after the FP sequence used in immunizations (AVGIGAVF).^1^ IgG elicited by Capped FP-Coil29 + Glyco_4_-Coil29 showed superior binding to FPv1 relative to all other treatment groups after 3 doses of FP immunogens and elicited detectable binding to FPv2 in some animals (Figure 4g).

After all doses of FP immunogens and SOSIP were administered, FPv1 and FPv2 binding were variable between animals, but serum from animals primed with capped nanofibers showed binding to both variants.

To measure the epitope specificity of endpoint sera, we utilized a previously described multiplexed binding assay to test antibody binding to a panel of epitopes from HIV-1 envelope protein.^26^ The overall profile of antibody binding to the panel of epitopes tested was significantly different for mice immunized with capped nanofibers than that elicited by FP-KLH (Figure 4h), indicating that this immunogen platform elicits a qualitatively unique priming environment for FP-focused immunization strategies. Taken together, the distinct FP binding antibody profiles generated by FP-Coil29 immunogens indicate the potential for improving FP-direct immunization approaches by diversifying the binding modes of FP-reactive antibodies.

### Designed nanofiber immunogens prime FP-specific antibodies in heterologous immunization of non-human primates

Because designed nanofiber immunogens displayed high lymph node accumulation, induced robust FP-directed antibody titers, and promoted IgG binding to multiple display formats of FP epitopes and heterologous mutants, we hypothesized that this formulation might be well-suited for preclinical development in higher order species. Mice are not capable of generating neutralizing responses to HIV-1 envelope and there are differences in the epitope specificity and functionality of envelope-reactive antibodies between rabbits and non-human primates.^27^ We conducted immunization studies in infant rhesus macaques as early life immunization is the most relevant use-case for potential HIV vaccines, since immunized infants have a long window of time for multidose prime-boost regimens and could be protected from infection prior to sexual debut.

We hypothesized that nanofiber-based immunogens might be capable of eliciting antibodies without adjuvant in non-human primates, as this effect has been observed robustly for nanomaterial-based immunogens administered in mice, and for Coil29 in particular.^15,28,29^ Because this was the first use of Coil29-based nanofiber immunogens in non-human primates, we wanted to determine the ability of these vaccines to generate antibodies without the need for additional adjuvants. For these reasons, we first tested the full formulation developed in mice (Capped FP-Coil29 + Glyco_4_-Coi29) in a non-human primate model wherein infant rhesus macaques were immunized without adjuvants 2-4 weeks after birth.

Three infant macaques received 5 doses of Capped FP-Coil29 + Glyco_4_-Coi29 followed by 2 doses of CH505 TF SOSIP. IgG responses to SOSIP, WT FP, and mutated variants of FP were measured throughout the course of the regimen (Figure 5a). FP nanofibers raised antibody responses to FP in all animals without the use of additional adjuvants. Unfortunately, anti-SOSIP IgG levels were negatively correlated with anti-FP titers (Figure 5b) and IgG binding to FPv2 was nearly abolished after 2 doses of SOSIP. This indicated that anti-FP responses were outcompeted by anti-SOSIP responses, presumably specific to other epitopes on SOSIP trimers. This may have occurred because FP-reactive B cells did not have sufficiently high affinity towards the full SOSIP trimer to compete with B cells which bound other epitopes on SOSIP trimers. Though we were encouraged that nanofiber immunogens did not require additional adjuvants to raise anti-FP titers in infant macaques, we sought to further optimize regimens to maintain FP-directed antibody responses after boosting with native-like SOSIP trimers.

**Figure 5:**
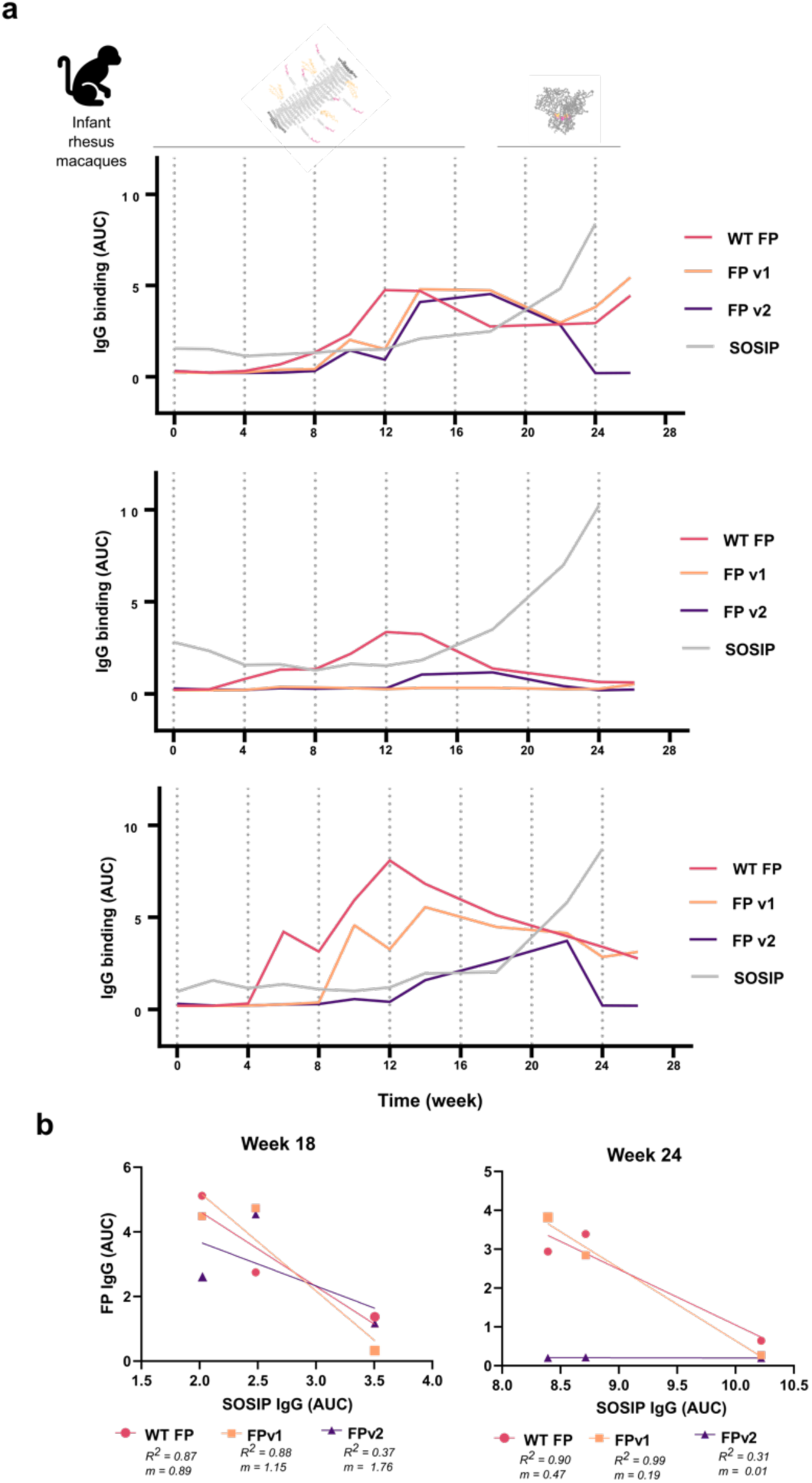
Designed immunogens raise FP-reactive antibodies in non-human primates without additional adjuvants. **(a)** Serum IgG binding to WT FP, SOSIP, FPv1 and FPv2 as measured by ELISA from biweekly blood collected from infant macaques immunized with 5 doses of Capped FP-Coil29 + Glyco_4_-Coil29 and 2 doses of CH505 TF SOSIP as indicated above. Responses to each ELISA probe are overlaid and each plot displays the responses of an individual animal. **(b)** Correlation of anti-SOSIP IgG with anti-FP IgG from the response curves in **a**.

We hypothesized that responses may not have been sufficiently potent without the inclusion of an adjuvant or may converge on non-native conformations of FP before SOSIP is introduced in the immunization scheme. For these reasons, we conducted a second study in infant rhesus macaques to test if varying the vaccination schedule and including an adjuvant might induce anti-FP responses which bound mutated FP variants and were boosted upon administration of SOSIP. In this study, new cohorts of infant macaques (5 animals per group) were given either sequential or alternating doses of Capped FP-Coil29 + Glyco_4_-Coi29 and BG505 SOSIP, each formulated with the adjuvant R848. We chose a small molecule adjuvant for this study to minimize alterations to the trafficking properties of FP nanofibers, as we expect could be the case for emulsion-based adjuvants or Alum. The two dosing regimens elicited distinct profiles in their induction of IgG directed towards FP, SOSIP, and mutated FP variants (Figure 6a).

**Figure 6:**
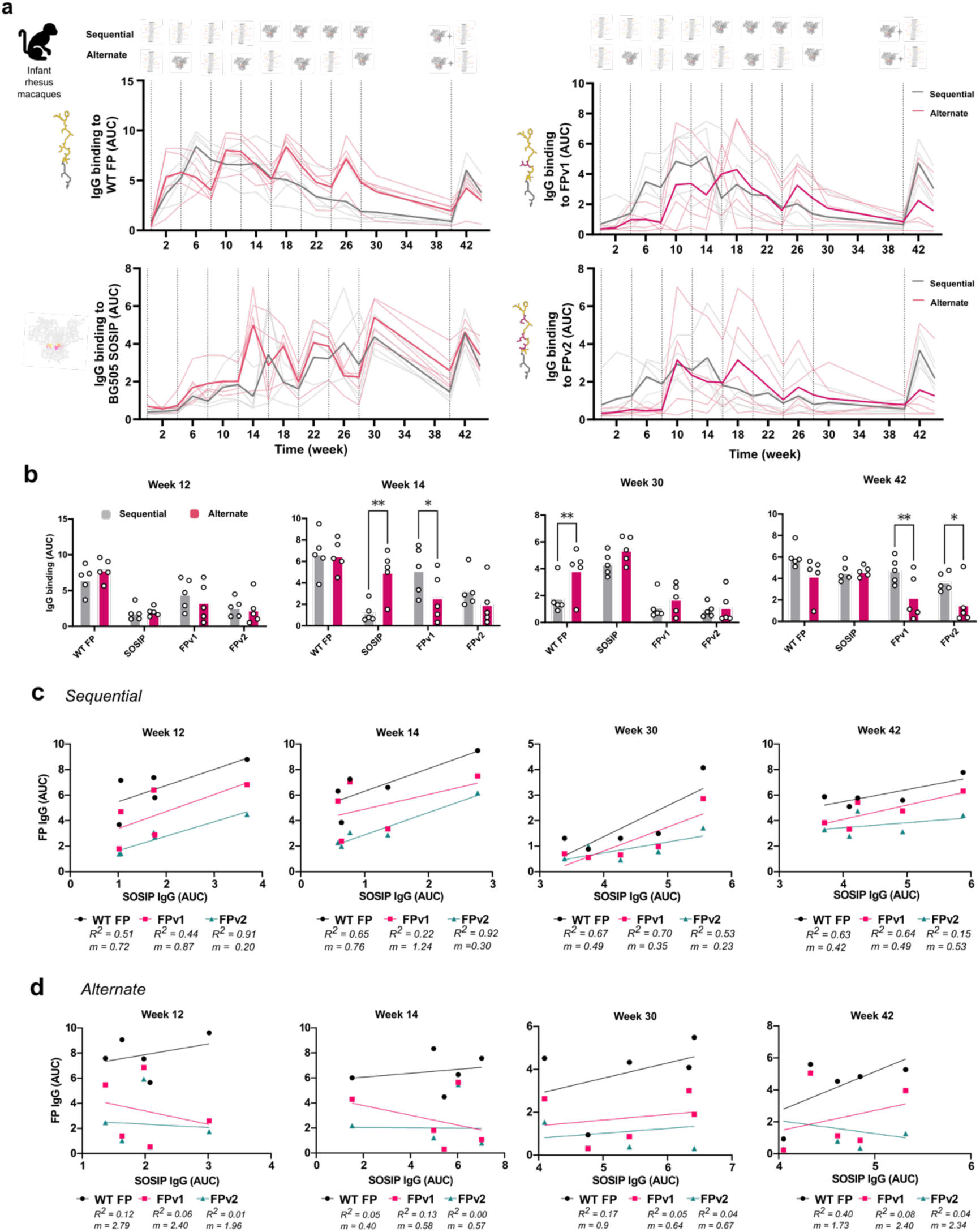
FP-reactive antibodies are primed by sequential immunization in non-human primates. **(a)** Serum IgG binding to WT FP, SOSIP, FPv1 and FPv2 as measured by ELISA from biweekly blood collected from infant macaques immunized with 4 doses of Capped FP-Coil29 + Glyco_4_-Coi29 and 4 doses of BG505 SOSIP each formulated with R848 according to the schedules indicated above. Mean responses are shown in bold with responses of each animal shown in thin lines. **(b)** Responses to the full panel of ELISA probes were compared at multiple timepoints using 2-way ANOVA and responses to each probe were compared using Fisher’s LSD test. **(c,d)** The relationship between anti-SOSIP IgG with anti-FP IgG from the time points shown above in **b**, R^2^ values and the slope of the fit for each correlation are shown to the right of the graph.

Early in the two immunization regimens (week 12), antibody levels for each antigen were not significantly different between sequential or alternating regimens, but after an additional dose of FP nanofibers (sequential) or SOSIP (alternating) at week 14, they diverged significantly. As expected, the alternating regimen yielded higher titers of SOSIP-reactive IgG (Figure 6b). Interestingly, WT FP-specific IgG levels remained indistinguishable between groups, but FPv1 titers were significantly higher in animals dosed with the sequential regimen than the alternating regimen. This suggests that the development of IgG which can bind multiple variants of FP requires repeated immunization with FP epitopes before exposure to full glycoprotein antigen.

Alternating SOSIP and FP nanofibers generated higher IgG binding to WT FP at week 30, possibly because the final dose of FP nanofibers was more recent for the alternating regimen than the sequential regimen. Responses were allowed to wane for 12 weeks prior to a final booster immunization of FP nanofibers and SOSIP (week 40). Interestingly, although alternating immunizations elicited higher anti-FP titers throughout the regimen, sequential FP-SOSIP dosing ultimately yielded a higher recall of IgG which bound FPv1 and FPv2 mutants. This suggests that repeated immunization with FP immunogens allows maturation of responses directed towards alternative sequences and/or conformations of FP without competition against other epitopes present on full HIV envelope trimers.

Over the course of the study, SOSIP-reactive antibody levels had a stronger and more positive correlation with FP-reactive antibodies for the sequential immunization regimen than the alternating regimen (Figure 6c,d). Across both immunization regimens and ELISA probes, IgG responses to FPv2 were most strongly correlated with SOSIP binding after 4 immunizations with FP nanofibers, prior to immunization with SOSIP trimers. This correlation suggests that the FP-reactive antibodies raised by nanofiber immunogens are capable of binding native-like envelope trimers and mutated variants of FP, which will be a critical step towards generating broadly reactive antibody responses in a clinical setting. After immunization with SOSIP, anti-FPv1 IgG levels were more strongly correlated with SOSIP-binding IgG than anti-FPv2 IgG levels, indicating that there may be some evolution of anti-FP responses toward a FPv1-reactive phenotype after administering response-maturing SOSIP immunizations.

Throughout the immunization regimen, FP-reactive antibodies were never strongly correlated with SOSIP-reactive antibodies for animals given alternating immunogens. This data suggests that the epitope specificity of SOSIP-binding IgG was widely spread in these animals, and the alternating regimen may not have been sufficient to focus immune responses on FP. In the future, alternating immunization regimens may be optimized with timing or adjuvant combinations to increase the formation of long-lived memory responses.

Using the sera from both of the aforementioned studies, we conducted pseudovirus neutralization assays but did not detect neutralization above background for any of the samples (Table S2, S3). Because these studies used infant macaques and different adjuvants than those tested in other FP-directed immunization strategies, it is possible they may elicit neutralizing antibodies in other contexts, or that individual antibody clones could be isolated and assessed for their neutralization potential.

Overall, these studies demonstrated that FP-specific antibodies can be elicited without adjuvants in non-human primates using the Coil29 vaccine platform. Sequential immunization with Capped FP-Coil29 + Glyco_4_-Coi29 was capable of priming FP-specific responses which were competitive against antibodies to other SOSIP epitopes, bound heterologous variants of FP, and were reactive to native-like HIV envelope SOSIP antigens. In future work, optimization of adjuvant, dosing, and immunization regimen schedules could be optimized to improve upon the responses we observed.

## Discussion

The investigation of FP-directed immunization highlights the utility of generating antibodies which are flexible to structural changes in the epitope of interest. In addition to FP-directed approaches for HIV-1, the ability to generate antibodies with a broad range of binding modes could improve the ability of vaccines to induce heterologous binding for other viruses. Widespread global diseases such as influenza and SARS-CoV2 may benefit from such approaches as these viruses are both widely circulating and present evolving variants over time.

Although neutralizing responses were not detected in this study, the ability to generate multiple antibody binding modes toward a known neutralizing epitope effectively allows for more “shots on goal” as the immune system generates antibodies with neutralization potential. In other studies, the angle at which antibodies approach neutralizing targets has influenced their neutralizing capacity significantly,^30,31^ and generating vaccines that diversify these possibilities could be broadly useful for vaccinating against viruses that are traditionally difficult to neutralize.

In non-human primates, we observed several interesting trends in the evolution of FP-reactive antibody responses across multiple immunization regimens. In future work, following the lineages of these antibodies over the course of heterologous immunization regimens may offer insight into the optimal strategy for maintaining breadth to mutated FP-strains while increasing affinity to native conformations of FP. Additionally, characterizing these unique binding profiles with tools such as polyclonal serum mapping with electron microscopy may indicate how the presentation of FP on nanofiber immunogens changes or diversifies antibodies towards this epitope.

The inclusion of branched glycomimetic peptides impacted the resultant trafficking and antibody responses toward peptide nanofibers, but the exact pathways which these peptides act on is not yet clear. Glycosylation has been shown to traffic nanomaterials to germinal centers via MBL-mediated complement activation^11^ which may be at play in this system. Additionally, the glycomimetic sequences selected for this study are known to bind Siglecs, which are widely expressed among immune cells. Specifically, CD169 is a known ligand of the peptide utilized here and is expressed on subcapsular sinus (SCS) macrophages, which transport materials to B cell follicles.^21^ If the peptides on these nanofibers bind SCS macrophages *in vivo*, their shuttling of materials to B cell follicles could improve germinal center reactions compared to materials transported via other pathways. We were surprised to note the strong immunogenicity of Capped FP-Coil29 without glycomimetic peptides although it did not show strong accumulation in dLNs to the same extent as the full formulation. This may indicate that other mechanisms are at play, such as changes in uptake, processing, or presentation which increase the immunogenicity of capped constructs regardless of the trafficking changes noted in Figure 2.

Glycomimetic peptides may have also influenced the conformational tolerance of elicited antibodies by altering the local structural context around displayed FP-epitopes. In addition to the trafficking benefits reported above, a brush-like network of disordered Glyco peptides may provide a similarly crowded environment to glycans on native HIV trimers, requiring binding antibodies to navigate masking hydrophilic contacts around the epitope of interest. If this is the case, selected B cells may have to adapt more diverse angles of approach to epitopes displayed in a variety of conformations in a shield of disordered Glyco peptides.

## Conclusions

The FP-Coil29 immunogens described here open up potential avenues for generating FP-directed antibody responses. Because of the diversified antibody binding modes elicited by Capped FP-Coil29 + Glyco_4_-Coil29 immunizations, these priming immunogens have significant promise for their ability to generate functional antibodies. Future generations of this approach could include mosaic assemblies displaying multiple strains of FP sequences and native-like antigens to simultaneously focus immune responses on FP epitopes while maturing responses towards native conformations. Interrogation of antibody binding modes generated by this display platform will be key in progressing the development of epitope-specific immunogens.

Self-assembling peptide vaccines are uniquely positioned to induce robust antibody responses to short peptide epitopes because of their opportunity for extremely dense epitope display, scalable synthetic production, and modular features.^32^ In comparison to KLH conjugates which have been explored in the context of FP-directed HIV immunization, peptide nanofibers offer advantages. Coil29 peptides can be produced entirely by chemical synthesis, which allows for scalable production and chemically well-defined therapeutics. Relative to KLH, these peptide-based systems have significantly improved batch-to-batch consistency because of their facile purification and modular composition. For deployment of an HIV-1 vaccine in limited resource settings, the need for cold-chain distribution presents significant logistical challenges. Peptide nanofibers are able to be stored as freeze dried powders^33^ which would provide immense benefit to the cost and timeliness of global vaccine distribution. In addition to practical advantages, nanofibers may allow for delivery across mucosal routes^33,34^, which could provide enhanced protection for vaginally or rectally transmitted HIV virus.^35–37^

Broadly speaking, the findings of this work show the potential for developing peptide-focused immunogens for heterologous prime-boost strategies. Because of their versatility in displaying a variety of peptide epitopes, this platform could be used to focus immune responses to fusion peptides or other known contiguous epitopes on antigenic surfaces.

## Acknowledgements

We thank Yaoying Wu for helpful discussions. We thank Jennifer Watanabe, Jodie Usachenko and the staff of Primate Resources and Logistics and Clinical Laboratories of the California National Primate Research Center for their expert assistance. We would like to acknowledge Kevin O. Saunders for providing CH505 TF SOSIP for studies in mice and infant macaques and John Moore for providing BG505 SOSIP for studies in infant macaques. This research was supported by the NIH (NIAID 1R01AI145016, 5R01AI167300) and the Duke Center for AIDS Research (CFAR). Awards P51OD011107 from the Office of Research Infrastructure Program, Office of The Director, National Institutes of Health, and U42OD010990, to the California National Primate Research Center. CNF and EJC were supported by the US National Science Foundation Graduate Research Fellowship (DGE-1644868). CNF was also supported by the Center for Biomolecular and Tissue Engineering NIH training grant (T32GM008555). MALDI was performed on an instrument supported by the North Carolina Biotechnology Center (2017-IDG-1018).

## Author Contributions

C.N.F. conceptualized studies, designed and synthesized immunogens, conducted immunization studies in mice, and prepared the manuscript. J.L.C. measured antibody responses from non-human primates, epitope breadth from mice, and analyzed data. N.L.V. designed and optimized trafficking experiments and IVIS measurements with E.J.C., R.K., C.N., S.S. and E.F.R B.C. conceptualized experiments and measured binding to F.P. variants with L.I. M.D. conducted ELISA binding assays from non-human primates. E.F.R. also coordinated shipment of immunizations for rhesus macaques. K.M.L. acquired circular dichroism measurements. K.V.R. designed experiments, immunized, and collected blood samples from rhesus macaques. S.P. and G.F. discussed findings and conceptualized experiments. J.H.C. designed immunogens, discussed findings, conceptualized experiments, acquired funding, and helped write the manuscript.

## Methods

### Peptide synthesis and purification

Peptides were synthesized by microwave-assisted solid phase synthesis with Fmoc-protected amino acids using a CEM Liberty Blue synthesizer, then cleaved and purified as previously described.^18^ To generate branched peptides, Fmoc-Lys(Fmoc)-OH residues were introduced to generate multiple reactive amines for chain lengthening. To extend peptides at multiple branching points, the stoichiometry of amino acids, Oxyma and DIC were doubled at each branching point to provide sufficient reagents for multiple chain extensions. After synthesis, branched peptides were cleaved and purified similarly to linear peptide variants. To generate KLH immunogens, FP-(EAAAK)_2_-C was coupled to maleimide activated Imject-KLH according to manufacturer instructions (Thermo Fischer).

### Nanofiber Formation and Atomic Force Microscopy

Nanofibers were formed by mixing 50% FP-Coil29 with 50% Coil29 or 10% Glyco_4_-Coil29 and 40% Coil29, depending on the formulation. Peptides are mixed as dry powders and dissolved in acetate buffer or 1.6 mM solutions of Coil29Caps in acetate buffer at 8mM total peptide concentration, as previously described.^18^ Prior to imaging or use for immunization, peptides were diluted to 2 mM in water and 10x PBS to yield a final solution of 1x PBS. Prior to AFM imaging, nanofibers are diluted 10-fold in ultrapure water and deposited on mica substrates, then rinsed with water and dried under a stream of nitrogen. AFM imaging was completed using a Bruker AFM operated in tapping mode with Bruker RTESPA-300 silicon tips.

### Viscometry of Nanofiber Assemblies

Nanofiber formulations were prepared identically to those described above for atomic force microscopy. The viscosity of nanofiber solutions was determined using a Malvern Kinexus rheometer with a 1mm plate. Frequency was ramped from 0.1-1000 Hz over 90 seconds, and sheer stress and viscosity were measured.

### Mouse studies and immunizations

To study immune responses to FP-Coil29 and FP-KLH immunogens, 8-week-old female C57BL/6 mice (Envigo) were injected subcutaneously at the tail base. For nanofiber immunizations (whether prime or boost), 2 separate 50 μL injections of 2mM total peptides were given. KLH was administered at a 10 μg dose to match previous studies, and CH505.TF SOSIP was given at a 10 μg dose after FP priming. Immunization schedules for each experiment are shown in the figure legends. For CpG adjuvanted immunizations, 10 μg of CpG was mixed with nanofibers, KLH or SOSIP prior to immunization. To analyze draining lymph node responses, inguinal lymph nodes were dissected and imaged by IVIS or processed for flow cytometry. For monitoring antibody responses, blood was collected from the submandibular vein at a biweekly frequency. All animal procedures were approved under IACUC protocol A264-18-11 at Duke University.

### IVIS Imaging of Lymph Nodes

To detect nanomaterial accumulation in lymph nodes, inguinal nodes were dissected post-injection and imaged in a Perkin Elmer Lumina III with an excitation laser at 640 nm and detection at 670 nm. Radiant efficiency in nodes was quantified using Living Image software and reported as the percentage of radiant efficiency of control lymph nodes harvested from naive mice.

### Intraperitoneal Uptake and Flow Cytometry

Linear Coil29 peptides were synthesized and fluorescently labeled with TAMRA at their N-terminus. Coil29 with a tetrameric branching point were synthesized as described above and subsequently labeled with Alexafluor647. To determine if linear and branched peptides could assemble in to the same nanofibers, nanofibers were prepared with either (1) linear peptides and branched peptides dissolved and assembled separately or (2) linear peptides and branched peptides mixed prior to solvation and assembly. Mice were then injected intraperitoneally with nanofibers and sacrificed after 4 hours. 2 mL of sterile PBS was injected in the peritoneal space, the abdomen was massaged, and peritoneal fluid was removed to harvest cells. Cells were then washed and stained for surface markers using the panel listed below. After being washed with PBS, cells were incubated with Live/Dead near-IR dye (Invitrogen) in 1:1000 dilution. Cells were then fixed with 2% formaldehyde in PBS and analyzed using a BD LSRII flow cytometer (BD Biosciences). Gating strategy to detect dendritic cells, B cells, and macrophages positive for one or both fluorophores is shown in Figure S2 (antibodies and vendors listed in Figure S2), representative plots shown in Figure S3.

### Flow Cytometry of Lymph Node Isolates

1-2 weeks after the final boost, inguinal lymph nodes were harvested from mice and crushed before filtering through a 70 μm cell strainer (VWR international LLC.) to break down the tissue, followed by purification using Lympholyte-M Cell Separation Media (Cedarlane). Lymphocyte density and viability were measured by MUSE Cell Analyzer (Sigma-Millipore). For B cell phenotyping, 2×10^6^ total calls were stained with fluorescently labeled antibodies IgG1 FITC, IgG2 FITC, IgG3 FITC, CD93 PE-CF954, IgM PE-Cy7, CD19 APC-R700, CD95 BV605, B220 BV650, CD138 BV711, CD23 BV786, and IgD BV510 (all from BD Biosciences), GL7 PE, CD21 PerCP-Cy5.5, CD11b BV570 (both from BioLegend), CD38 PE-Cy5 (eBioscience). After being washed with PBS, cells were incubated with Live/Dead near-IR dye (Invitrogen) in 1:1000 dilution. Cells were then fixed with 2% formaldehyde in PBS and analyzed using a BD LSRII flow cytometer (BD Biosciences). Analysis of specific cell populations was executed following the gating schemes shown in Figure S4.

### ELISPOT Detection of Antigen-Specific T cells

ELISPOT was used to detect peptide-responsive T cells as previously described.^16^ Briefly, spleens were harvested from mice 1-2 weeks after the final immunization. Spleens were crushed and filtered through a 70 μm cell strainer and washed in cold PBS with 2% FBS. Red blood cells were then lysed with ACK lysing buffer, cells were washed, and lymphocytes were isolated using Lympholyte M. 2.5×10^6^ cells were then mixed with stimulating peptides (5 μM) and cultured on membrane plates coated in IL-4 or IFNγ capture antibodies for 48 hours. After culture, cells were removed, and plates were washed in PBST before adding the corresponding detection antibodies to IL-4 and IFNγ plates for 2 hours at room temperature. Plates were then washed again and treated with streptavidin-ALP before a final wash and treatment with SigmaFast substrate. Plates were then monitored for the appearance of spots and washed with ultrapure water to stop spot formation. Spot counts were quantified by ZellNet Consulting for analysis.

### ELISA probes

FP_C_ was synthesized by first synthesizing the FP sequence (AVGIGAVFL) on resin as described above, followed by an SGSG linker. Before cleaving the peptide from resin, a biotin group was added, yielding biotin-SGSG-AVGIGAVFL. FP_N_, FPv1, and FPv2 were synthesized so that the N-terminus of FP would be unmodified for binding in ELISA, analogous to how it is displayed on nanofiber immunogens. A helical spacer and single cysteine residue were added C-terminally from the FP epitope, yielding the full sequence AVGIGAVFL-SGSG-(EAAAK)_2_-SGSG-C. After cleavage from resin, a biotin group was added using a maleimide-(PEG)_2_-biotin linker (Thermo Fisher). Adsorbed FP was synthesized with the antigenic sequence alone (AVGIGAVFL).

### ELISA Analysis of Serum Antibodies

For FP_N_ and FP_C_ ELISAs, Corning 96-well high binding plates were coated overnight with streptavidin (2 μg/mL) before washing and applying biotinylated FP peptides (all sequences listed in **Error! Reference source not found.**) at 20 μg/mL or PBS to alternating columns and incubating for 1 hour. After washing, plates were blocked with SuperBlock and murine serum diluted in 1% BSA in PBST was applied to peptide-coated and uncoated wells at dilutions spanning 1:100 to 1:10,000 for 2 hours. After serum incubation, plates were washed, and murine-IgG HRP antibody was applied for 45 minutes. Finally, plates were washed, and bound antibody was measured using TMB and a phosphoric acid stop solution.

For FP_p_ ELISAs, the same method described above was followed, except that FP peptides dissolved in PBS were coated overnight at 20 μg/mL before washing and blocking the following day.

To detect SOSIP-binding antibodies, plates were coated overnight in PGT145 antibody (4 μg/mL), washed, and blocked with SuperBlock for 1 hour. SOSIP was then applied (10 μg/mL) and incubated for 2 hours at room temperature. After washing, murine serum diluted in 1% BSA in PBST was applied to wells at dilutions spanning 1:100 to 1:10,000 for 2 hours. After serum incubation, plates were washed, and murine-IgG HRP antibody was applied for 45 minutes. For all ELISAs, plates were washed, and bound antibody was measured using TMB and a phosphoric acid stop solution. To detect antibody binding, plates were read using a SpectraMax plate reader (450 nm).

### Multiplexed antigen binding assay (BAMA)

Multiplexed antigen binding experiments were conducted as previously described.^26,38^ Briefly, a panel of HIV antigens including 1086C. gp120,^39^ MN. gp120 gDneg (clade B),^40^ A244 gp120 gDneg (Clade AE),^40^ A1.con.env03 gp140 (consensus clade A Env gp140 with deletions in the gp41 cleavage site and fusion domain),^41^ and B.con.env03 gp140 (consensus clade B Env gp140)^40^ were conjugated to beads. Mouse sera were diluted in diluent (1% skim milk, 5% normal goat serum, 0.05% tween 20 in 1X PBS; pH 7.4) to generate six 6-fold serially diluted samples starting from 1:10 dilution. Antigen coated beads were then incubated with sera at room temperature for 30 min and washed 3 times (0.1% BSA, 0.02% Tween 20, 0.05% sodium azide in 1X PBS; pH 7.4). PE-conjugated goat anti-Mouse IgG (SouthernBiotech) (2 μg/ml) was then added at room temperature for 30 min, washed 3 times, and then measured by a Bio-Plex 200 System. Results were expressed as mean fluorescence intensity (MFI).

### Immunization studies in rhesus macaques

Infant rhesus macaques (*Macaca mulatta*) of Indian origin were from the breeding colony of the California National Primate Research Center, which is negative for type D retrovirus, simian immunodeficiency virus, simian lymphocyte tropic virus type 1. Infant macaques were reared by their mothers until they were weaned at approximately 6 months of age. All procedures are done according to established standard operating procedures. Animal care followed the *Guide for Care and Use of Laboratory Animals* by the Institute for Laboratory Animal Research. Animal procedures were approved by the UC Davis Institutional Animal Care and Use Committee prior to study initiation. For immunizations and sample collections, animals were sedated with ketamine HCl (5-30 mg/kg IM). EDTA-anticoagulated blood samples were collected via venipuncture at defined time points and processed for plasma

For the first study (Figure 5), Infants were given a priming immunization within 4 weeks after birth. Capped FP-Coil29 + Glyco_4_-Coil29 nanofibers were prepared as described above, then diluted in sterile PBS to 0.2mM prior to injection. A 500 μL suspension of nanofibers were injected subcutaneously without any additional adjuvants on the upper back of the animals. Immunizations were administered every 4 weeks and blood was collected every 2 weeks. Macaques received a total of 5 doses of nanofibers, followed by 2 doses of CH505 SOSIP. SOSIP was mixed with 3M-052 in equal volumes immediately prior to injection. 3 animals were immunized in this study, and responses were measured and reported independently.

For the second study (Figure 6), 8 immunizations were administered 4 weeks apart. Sequentially immunized animals received 4 doses of nanofibers, followed by 4 doses of BG505 SOSIP. Alternating immunizations administered 4 doses of nanofibers and 4 doses of BG505 SOSIP, alternating immunogens as shown in Figure 6. Nanofibers were administered at 0.2 mM with 100 μg of R848. SOSIP was administered at a 50 μg dose, also formulated with 100 μg of R848. A final dose of 60 μg of SOSIP and 700 uL of 0.2 mM nanofibers was administered at week 40. Over the course of the entire regimen, blood was collected every 2 weeks. All solutions were verified to have an endotoxin level <0.01 EU prior to administration. R848 was purchased from InvivoGen (tlrl-r848-1) and was mixed with nanofibers or SOSIP immediately prior to immunization.

### Serum Neutralization

Neutralizing antibody titers were determined using the standardized TZM-bl cell assay as previously described (PMCID: PMC4040342). To perform the assay, plasma samples were serially diluted threefold, ranging from 1:20 to 1:43,740, and then mixed with the target HIV pseudovirus. The mixture was incubated at 37°C with 5% CO₂ for 1 to 1.5 hours to allow neutralization to occur. Following this step, O-(diethylaminoethyl)–dextran (10 μg/ml; Sigma-Aldrich, #D9885) and 1 × 10⁵ TZM-bl cells (NIH AIDS Reagent Program, #8129) were introduced into each well. After 48 hours of incubation, 150 μl of supernatant was removed, and 100 μl of Bright-Glo luciferase substrate (Promega, #E264CX) was added to each well. The resulting mixture (150 μl) was then transferred to Corning 96-well flat-bottom black plates (VWR, #29444-018), and luminescence was quantified using a Victor X3 Multilabel Plate Reader (PerkinElmer).

