## Supplemental Information for "Designed nanofiber immunogens prime antibody responses to diverse conformations of the HIV-1 fusion peptide"

### Supplementary Text

We have previously noted that different peptide epitopes can influence the length of assembled nanofibers. Appending particularly hydrophobic epitopes such as SIINFEKL to Coil29 has resulted in shorter nanofibers than those bearing more hydrophilic peptides.[1] In previous work, Coil29Caps shortened nanofiber assemblies, but in this study, uncapped nanofibers bearing FP we already short, aggregated, and poorly assembled. In this study, the inclusion of Coil29Caps resulted in longer assemblies than nanofibers assembled without Coil29Caps. We hypothesize that the stabilization of exposed ends of Coil29 nanofibers limited the possible association of FP epitopes with Coil29, enabling longer nanofiber assemblies which were difficult to assemble otherwise. We also noted that the inclusion of Glyco$_4$-Coil29 in assemblies increased nanofiber length and decreased the viscosity of bulk nanofiber solutions. We interpreted this change in viscosity as an indication that both branched Glyco$_4$-Coil29 peptides and capping peptides are necessary to generate a solution of nanofibers which do not further associate into an extended network. This may be attributable to the existence of brush-like conformations of branched peptides acting to repel nanofibers from each other. However, this phenomenon only appears to be at play when nanofibers are sufficiently stabilized at their tips by capping peptides, allowing stable nanofibers to form.

### Table S1: FP-Coil29 Sequences

| | |
|---|---|
| FP-Coil29 | AVGIGAVFL-SGSG-(EAAAK)$_2$-SGSG QARILEADAEILRAYARILEAHAEILRAQ-$NH_2$ |
| Alexa$^{647}$-Coil29 | AlexaFluor$^{647}$-SGSG-(EAAAK)$_2$-SGSG QARILEADAEILRAYARILEAHAEILRAQ-$NH_2$ |
| Coil29 | Ac-QARILEADAEILRAYARILEAHAEILRAQ-$NH_2$ |
| Coil29Cap a/d | Ac-QSRITEADSEITRAYSRITEAHSEITRAQ-$NH_2$ |
| Coil29Cap c/f | Ac-QARTLESDAETLRSYARTLESHAETLRSQ-$NH_2$ |
| Biotin-FP | Biotin-SGSG-AVGIGAVFL |
| FP-Biotin | AVGIGAVFL-SGSG-(EAAAK)$_2$-SGSG-C-PEG-biotin |
| FP | AVGIGAVFL |

PEG = polyethylene glycol, FP epitope, Coil29 assembling sequence, Ac = acetyl

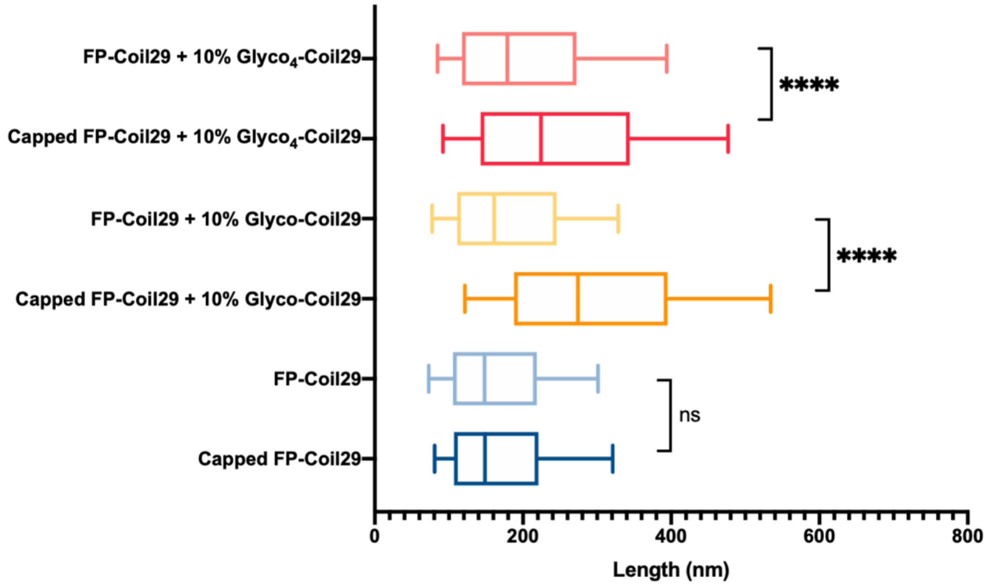


**Figure S1: Length of FP nanofibers. FP nanofibers imaged by AFM were measured using ImageJ and lengths were compared using One-way ANOVA. Capped and Uncapped formulations were compared using Bonferroni post-hoc testing.****p<0.0001**

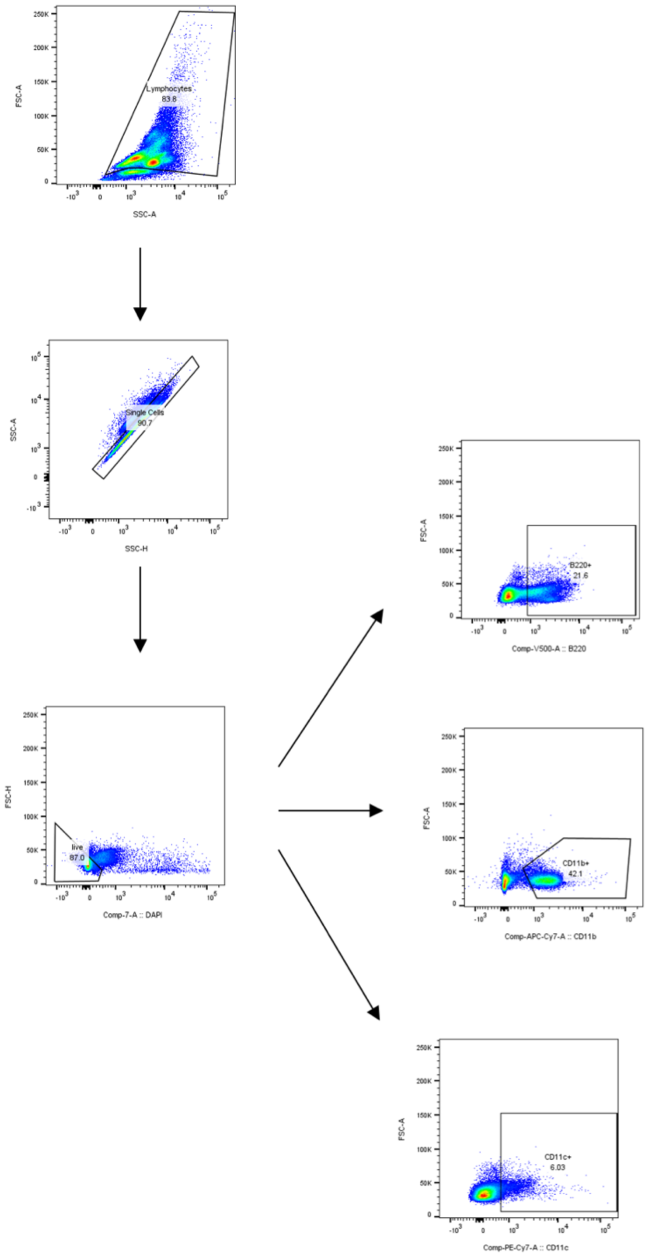


| Channels | Markers | Clones | Vendor |
|---|---|---|---|
| FITC | CD11c | N418 | Biolegend |
| PerCP/PerCP-Cy5.5 | F4/80 | BM8 | Biolegend |
| PE-Cy7 | MHCII | M5/114.15.2 | Biolegend |
| APC-Cy7 | CD11b | M1/70 | BD Biosciences |
| BV510 | B220 | RA3-6B2 | BD Biosciences |

**Figure S2. Gating of immune subsets from intraperitoneal lavage**

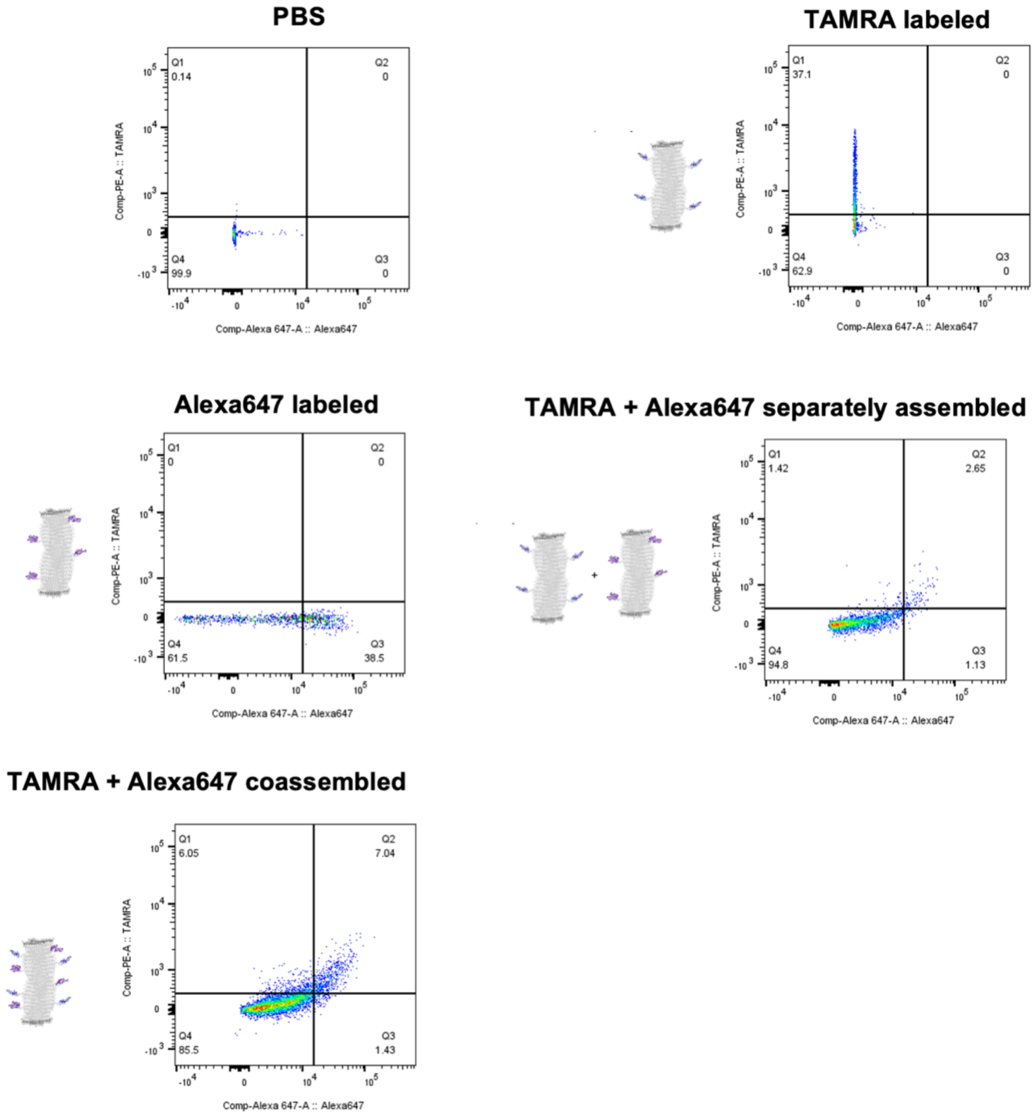


**Figure S3. Representative flow plots of Alexa647 and TAMRA in gated populations of cells harvested after intraperitoneal injection and lavage**

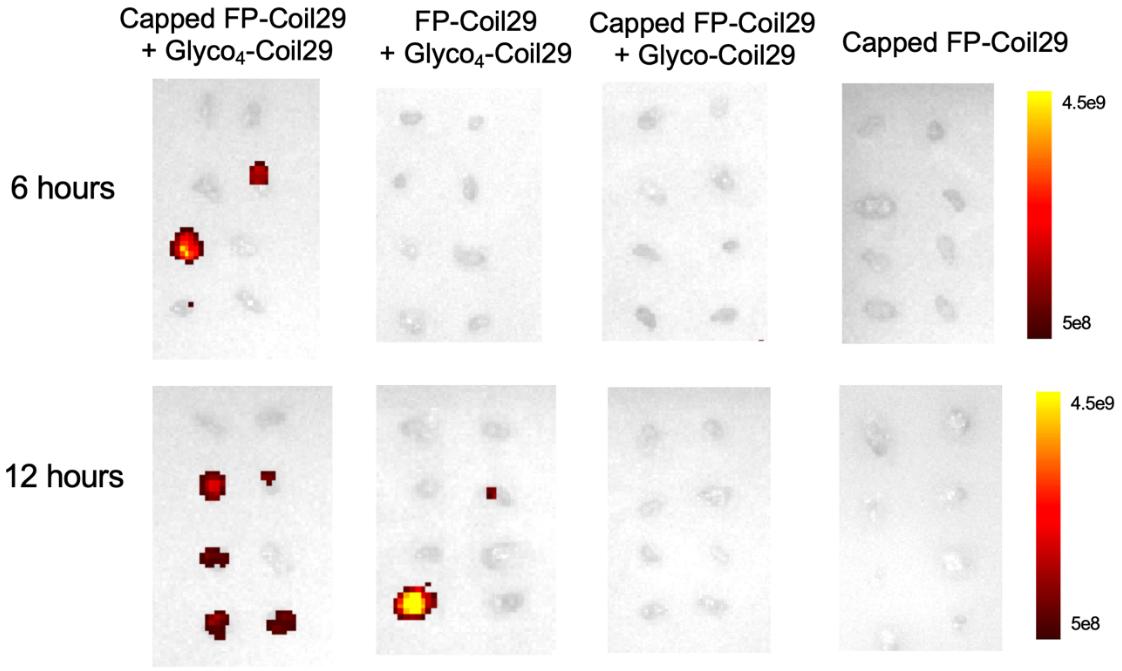


**Figure S4. IVIS images of excised lymph nodes at 6- and 12-hours post injection**

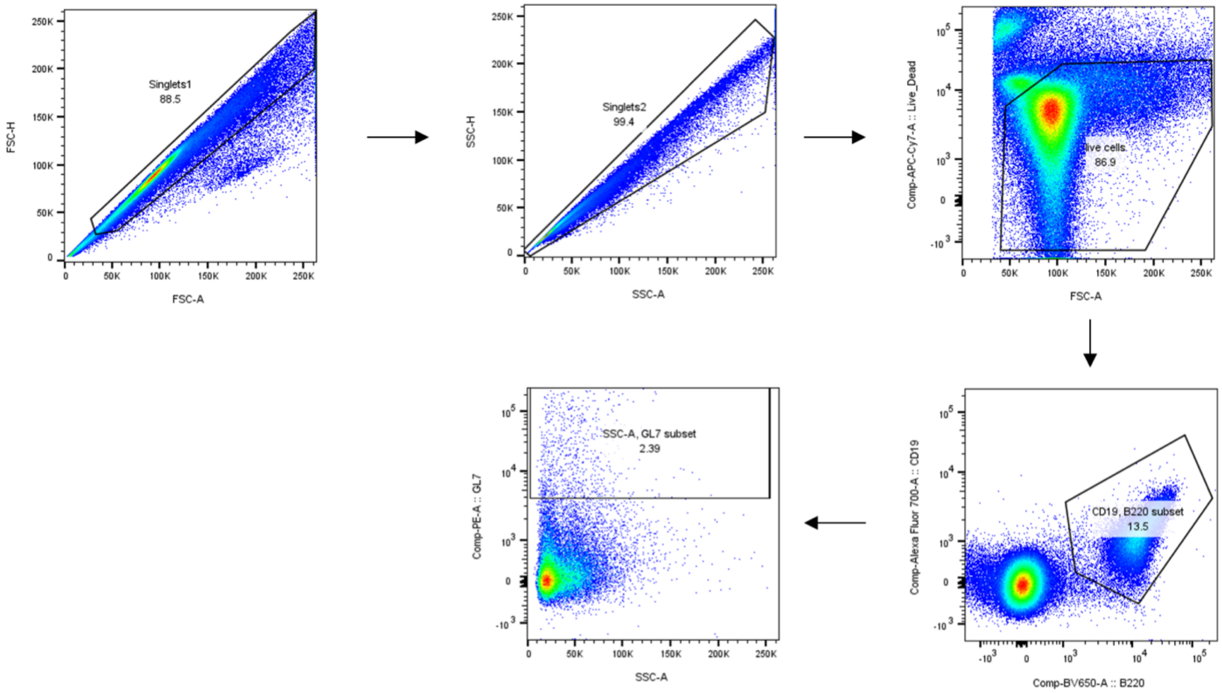


**Figure S5: Gating of GL7hi germinal center B cells**

| Animal ID | Vaccine | Week | Env | $ID_{50}$ |
|---|---|---|---|---|
| 49872 | FP-Coil29 | 6 | CH505 w4.3 | <20 |
| 49890 | FP-Coil29 | 6 | CH505 w4.3 | <20 |
| 49922 | FP-Coil29 | 6 | CH505 w4.3 | <20 |
| 49872 | FP-Coil29 | 10 | CH505 w4.3 | <20 |
| 49890 | FP-Coil29 | 10 | CH505 w4.3 | <20 |
| 49922 | FP-Coil29 | 10 | CH505 w4.3 | <20 |
| 49872 | FP-Coil29 | 14 | CH505 w4.3 | <20 |
| 49890 | FP-Coil29 | 14 | CH505 w4.3 | <20 |
| 49922 | FP-Coil29 | 14 | CH505 w4.3 | <20 |
| 49872 | FP-Coil29 | 30 | CH505 T/F | <20 |
| 49890 | FP-Coil29 | 30 | CH505 T/F | <20 |
| 49922 | FP-Coil29 | 30 | CH505 T/F | <20 |
| 49872 | FP-Coil29 | 34 | CH505 T/F | 24 |
| 49890 | FP-Coil29 | 34 | CH505 T/F | 31 |
| 49922 | FP-Coil29 | 34 | CH505 T/F | <20 |
| 49872 | FP-Coil29 | 18 | CH505 w4.3 N611A | <20 |
| 49890 | FP-Coil29 | 18 | CH505 w4.3 N611A | <20 |
| 49922 | FP-Coil29 | 18 | CH505 w4.3 N611A | 25 |
| 49872 | FP-Coil29 | 22 | CH505 w4.3 N611A | <20 |
| 49890 | FP-Coil29 | 22 | CH505 w4.3 N611A | <20 |
| 49922 | FP-Coil29 | 22 | CH505 w4.3 N611A | <20 |
| 49872 | FP-Coil29 | 26 | CH505 w4.3 N611A | <20 |
| 49890 | FP-Coil29 | 26 | CH505 w4.3 N611A | <20 |
| 49922 | FP-Coil29 | 26 | CH505 w4.3 N611A | 32 |
| 49872 | FP-Coil29 | 30 | CH505 w4.3 N611A | 22 |
| 49890 | FP-Coil29 | 30 | CH505 w4.3 N611A | 85 |
| 49922 | FP-Coil29 | 30 | CH505 w4.3 N611A | 45 |
| 49872 | FP-Coil29 | 34 | CH505 w4.3 N611A | 53 |
| 49890 | FP-Coil29 | 34 | CH505 w4.3 N611A | 189 |
| 49922 | FP-Coil29 | 34 | CH505 w4.3 N611A | 40 |

**Table S2: Serum neutralization $ID_{50}$ from NHP immunized with unadjuvanted FP nanofibers (binding data shown in Figure 5)**

| Animal ID | Week | Experimental Group | BG505 332N HIV | BG505 332N N611A |
|---|---|---|---|---|
| 50228 | 30 | FP-coil29/SOSIP sequential | <20 | <20 |
| 50424 | 30 | FP-coil29/SOSIP sequential | <20 | <20 |
| 50599 | 30 | FP-coil29/SOSIP sequential | <20 | <20 |
| 50652 | 30 | FP-coil29/SOSIP sequential | <20 | <20 |
| 50702 | 30 | FP-coil29/SOSIP sequential | <20 | <20 |
| 50236 | 30 | FP-coil29/SOSIP alternate | <20 | <20 |
| 50593 | 30 | FP-coil29/SOSIP alternate | <20 | <20 |
| 50610 | 30 | FP-coil29/SOSIP alternate | <20 | <20 |
| 50653 | 30 | FP-coil29/SOSIP alternate | <20 | <20 |
| 50709 | 30 | FP-coil29/SOSIP alternate | <20 | <20 |
| 50228 | 42 | FP-coil29/SOSIP sequential | <20 | <20 |
| 50424 | 42 | FP-coil29/SOSIP sequential | <20 | <20 |
| 50599 | 42 | FP-coil29/SOSIP sequential | 36 | 28 |
| 50652 | 42 | FP-coil29/SOSIP sequential | <20 | <20 |
| 50702 | 42 | FP-coil29/SOSIP sequential | <20 | <20 |
| 50236 | 42 | FP-coil29/SOSIP alternate | <20 | <20 |
| 50593 | 42 | FP-coil29/SOSIP alternate | <20 | <20 |
| 50610 | 42 | FP-coil29/SOSIP alternate | <20 | 28 |
| 50653 | 42 | FP-coil29/SOSIP alternate | <20 | <20 |
| 50709 | 42 | FP-coil29/SOSIP alternate | <20 | <20 |

**Table S3: Serum neutralization $ID_{50}$ from NHP immunized with sequential or alternating doses of FP nanofibers (binding data shown in Figure 6)**
